# Investigating the effectiveness of school health services delivered by a health provider: a systematic review of systematic reviews

**DOI:** 10.1101/543868

**Authors:** Julia Levinson, Kid Kohl, Valentina Baltag, David Ross

## Abstract

Schools are the only institution regularly reaching the majority of school-age children and adolescents across the globe. Although at least 102 countries have school health services, there is no rigorous, evidence-based guidance on which school health services are effective and should be implemented in schools. To investigate the effectiveness of school health services for improving the health of school-age children and adolescents, a systematic review of systematic reviews (overview) was conducted. Five databases were searched through June 2018. Systematic reviews of intervention studies that evaluated school-based or school-linked health services delivered by a health provider were included. Review quality was assessed using a modified Ballard and Montgomery four-item checklist. 1654 references were screened and 20 systematic reviews containing 270 primary studies were assessed narratively. Interventions with evidence for effectiveness addressed autism, depression, anxiety, obesity, dental caries, visual acuity, asthma, and sleep. No review evaluated the effectiveness of a multi-component school health services intervention addressing multiple health areas. Strongest evidence supports implementation of anxiety prevention programs, indicated asthma education, and vision screening with provision of free spectacles. Additional systematic reviews are needed that analyze the effectiveness of comprehensive school health services, and specific services for under-researched health areas relevant for this population.

## INTRODUCTION

The World Health Organization (WHO) launched the Global School Health Initiative in 1995 with the goal to improve child, adolescent and community health through health promotion and programming in schools [1]. This initiative is dedicated to promoting development of school health programs and increasing the number of health-promoting schools, characterized by WHO as “a school constantly strengthening its capacity as a healthy setting for living, learning and working” [1]. In 2000, WHO, the United Nations Educational, Scientific and Cultural Organization (UNESCO), the United Nations Children's Fund (UNICEF) and the World Bank developed a partnership for Focusing Resources on Effective School Health – a FRESH Start approach [2]. The FRESH framework promotes four pillars: health-related school policies, provision of safe water and sanitation, skills-based health education and school-based health and nutrition services [2]. While various guidance documents have been published by United Nations (UN) organizations addressing a range of services from oral health to malaria [3–7], there is no internationally accepted guideline regarding school health services. This systematic review of systematic reviews, henceforth referred to as an overview, will inform the upcoming development of a WHO guideline that addresses one pillar of the FRESH framework: school health services delivered by a health provider.

Schools offer a unique platform for health care delivery. In 2015, the global means for the primary and secondary net school enrollment rates were 90% and 65%, respectively, thus the potential reach of school health services is wide [8]. Additionally, a recent review found that school-based or school-linked health services already exist in at least 102 countries [9]. The 2017 Global Accelerated Action for the Health of Adolescents (AA-HA!) implementation guidance calls for the prioritization of school health programs as an important step towards universal health coverage and urges that “Every school should be a health promoting school” [10].

The primary objective of this overview was to explore the effectiveness of school-based or school-linked health services delivered by a health provider for improving the health of school-age children and adolescents. Through a comprehensive literature search, the overview aimed to identify health areas and specific school health service interventions that have at least some evidence of effectiveness. It was also designed to suggest further research in areas where recent systematic reviews (SRs) exist, but with insufficient evidence. Finally, the overview aimed to identify the health areas and specific school health services interventions for which no SRs were found, whether because the primary literature does not exist or where there are primary studies but no SR has been conducted.

## METHODS

This overview was conducted using the preferred reporting items for systematic reviews and meta-analyses (PRISMA) [11]. A protocol was developed *a priori* that outlined the overview objectives, aims, operational definitions, search strategy, inclusion/exclusion criteria, and quality appraisal methods. This document was followed throughout the review process and is available in S1 Appendix.

### Search strategy

PubMed, Web of Science, ERIC, PsycINFO, and the Cochrane Library were searched systematically. A detailed search strategy was iteratively developed in consultation with a librarian experienced in SRs and an expert in school health services. The search strategy was developed for PubMed and then adapted for the other four databases. The search strategy is presented in S2 Appendix. Searches were performed on June 15, 2018. Any existing overviews or SRs of SRs that emerged from the searches were not themselves included, but the SRs within them were extracted and screened. Additionally, reference lists of included articles were scanned for any relevant SRs.

### Eligibility criteria

SRs were included in this overview if at least 50% of the studies within the SR fulfilled the following criteria: (a) participants were children (ages 5-9) or adolescents (ages 10-19) enrolled in schools; (b) interventions were school-based or school-linked health services, involved a health provider (see definitions in S1 Appendix), and were of any duration or length of follow-up; (c) intervention effectiveness was compared to either no intervention, an alternative intervention, the same intervention in a different setting (i.e. not in schools), an active control, or a waitlist control; (d) interventions aimed to improve some aspect of health; and (e) study designs were either randomized controlled trials (RCTs), quasi-experimental studies (QEs), or other non-randomized intervention studies. There were no date restrictions on publication of included SRs. In addition to these criteria for included studies, the SRs themselves had to fulfill the following criteria: (a) included the words “systematic review” in the title or abstract; (b) outlined inclusion criteria within the methods section; (c) published in peer-reviewed journals and indexed before June 15, 2018; (d) published in the English language. In addition to SRs that did not meet these inclusion criteria, SRs were excluded if the review was superseded by a newer version.

### Study selection

Citations identified from the systematic search were uploaded to Covidence systematic review software [12] and duplicates were automatically deleted. Two reviewers (KK and JL) screened all titles and abstracts using the inclusion/exclusion criteria and excluded all articles that were definitely ineligible. Articles that received conflicting votes (ineligible vs. potentially or probably eligible) were discussed and consensus was reached. The same two reviewers screened the full text of all the potentially or probably eligible articles using a ranked list of the inclusion criteria (S1 Appendix). Reasons for exclusion were selected from the ranked list. If consensus was not possible during title/abstract or full text screening, a third reviewer (DR), who had the casting vote, would have been asked to independently screen the article. This was never required as consensus was always reached.

### Data collection

One reviewer (JL) extracted summary data from each selected article using a customized standard form with independent data extraction performed for 15% of included SRs by one of the other reviewers (DR or KK). There was 92% agreement between reviewers for all items within the standard form, with discrepancies only in level of detail. Data items included the research design of the SR and primary studies, sample description and setting, intervention characteristics, outcomes, meta-analysis results, quality appraisal, and conclusions.

### Synthesis of results

Due to the heterogeneity of the SRs included in this overview, it was not possible to perform a meta-analysis. Outcome measures were collected from included studies.

### Risk of bias

Risk of bias within primary studies was recorded in S4 Appendix. Risk of bias across SRs was determined using Ballard and Montgomery’s four-item checklist for overviews of SRs [13]. These items include: (1) overlap (see below), (2) rating of confidence from the AMSTAR 2 checklist [14], (3) date of publication, and (4) match between the scope of the included SRs and the overview itself.

An important consideration in overviews is the degree of overlap, or the use of the same primary study in multiple included SRs. High overlap can contribute to biased results [15]. This overview used the corrected covered area (CCA), a comprehensive and validated measure, to determine overlap [16]. The CCA is calculated using three variables: the number of “index” publications (r), the number of total publications (N), and the number of SRs within the overview (c). An “index” publication is the first appearance of a primary study within an overview. The formula for the CCA is:

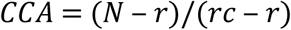

CCAs can be interpreted as indicating slight, moderate, high or very high overlap with scores of 0-5, 6-10, 11-15, or >15, respectively [16].

The AMSTAR 2 checklist [17] was used to appraise quality of included SRs. One reviewer (JL) assessed all SRs and a second (KK) duplicated appraisal of 10%, with 94% agreement and only minor disagreements that did not impact grades of confidence. Following the recommendation of the AMSTAR 2 developers [17], individual ratings were not combined into an overall score. Instead, the authors determined which of the 16 items on the checklist were critical for this overview and which of the items were non-critical. Building on a method suggested by Shea and colleagues [17], grades of confidence in the results of each SR were generated based on critical flaws and non-critical weaknesses. The grading system is available in S6 Appendix. Confidence in results ranged from high (three or fewer non-critical weaknesses) to critically low (more than three critical flaws with or without non-critical weaknesses) [17].

The final two items in the four-item checklist are (3) date of publication to ensure that results are up-to-date, and (4) matching of the scope to confirm that primary studies within SRs are relevant to the overview. This overview considered SRs published in 2016, 2017, or 2018 to be up-to-date. Although all SRs were included if at least 50% of the primary studies within them fulfilled all inclusion criteria, SRs where closer to 100% of primary studies fulfilled all criteria were considered to be better matched than those with closer to 50%.

## RESULTS

### Results of the search

1720 references were identified from the systematic literature search. 570 were duplicates, leaving 1150 titles and abstracts to be screened. After removal of 705 articles with definitely ineligible titles/abstracts, 445 full text articles were screened using the ranked inclusion/exclusion criteria (S1 Appendix). 425 of these articles were ineligible. The most common reasons for exclusion were, first, if at least 50% of studies included within a SR were unclear for at least one criterion (e.g. unclear if a health provider was involved), followed by if less than 50% of studies within a SR fulfilled all inclusion criteria. These reasons for exclusion applied to 183 and 126 SRs, respectively. The full list of the specific reasons for exclusion of full texts is available in S3 Appendix. Figure 1 displays the PRISMA flowchart for the search.

**Fig 1.**
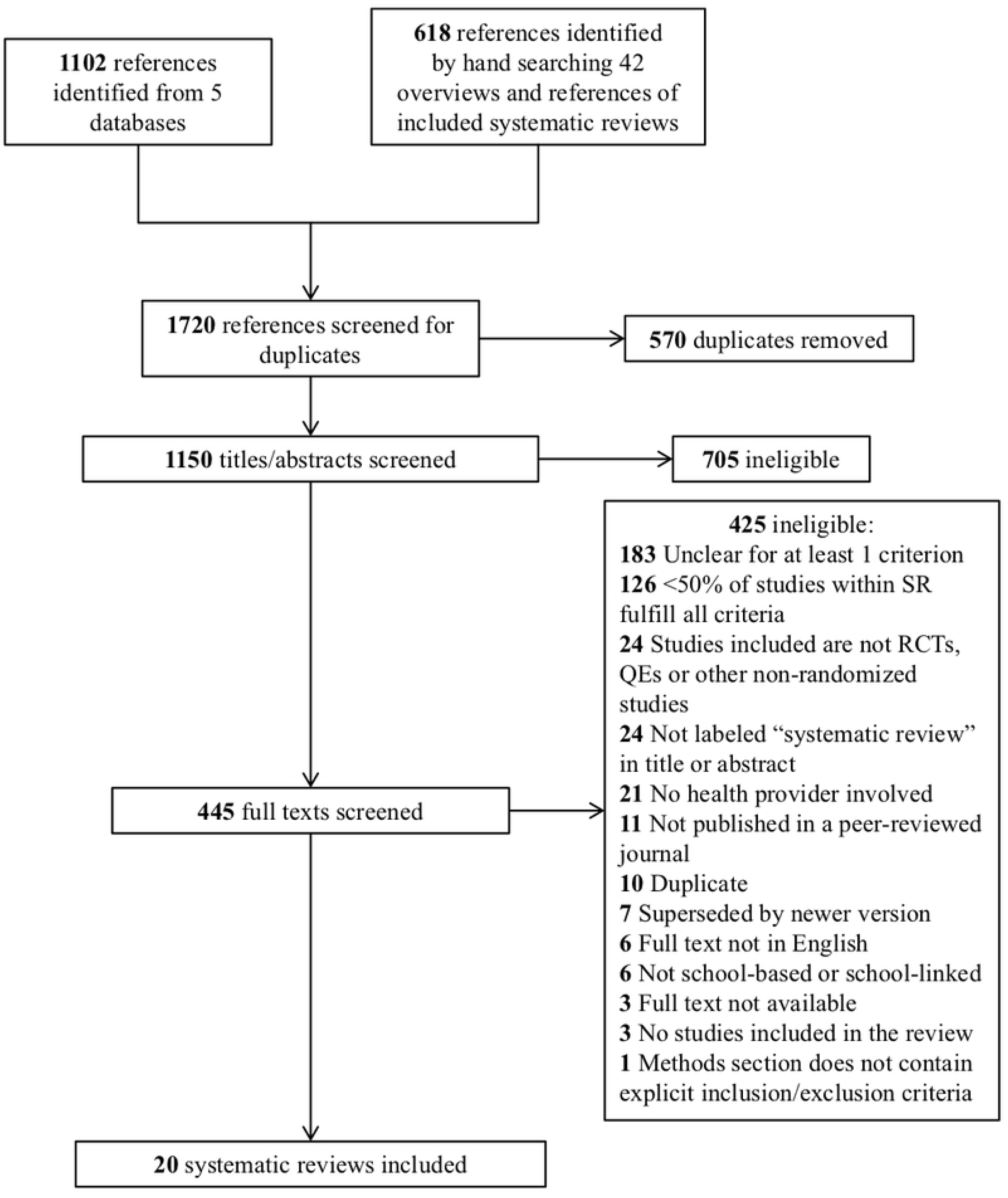
PRISMA flowchart showing reasons for exclusion of potential systematic reviews

### Characteristics of included systematic reviews

20 SRs fulfilled all inclusion criteria and were included in this overview. These SRs contained 270 primary studies, 225 of which were unique and 45 were included in more than one SR. The SRs were written in English and published between 2006 and 2018. Primary studies included within the SRs were published between 1967 and 2017. Eleven of the SRs used meta-analysis to combine results [18–28], whereas the remaining nine SRs narratively synthesized results [29–37]. Eleven SRs included studies located in countries with high-income or upper-middle income economies only [19,20,23–25,29–33,37,38,39]. Six SRs included at least one study from countries with lower-middle income or lower income economies [18,21,22,28,35,36]. The final three SRs either did not state the locations of included studies [27,34] or provided regions rather than specific country locations [26].

To be included in this overview, at least 50% of studies within each SR had to fulfill all inclusion criteria. In four SRs, 100% of included studies fulfilled all inclusion criteria [18,27,29,33], although Brendel and colleagues [29] only included one study in total. In another four SRs, 75% to 99% of included studies fulfilled all the inclusion criteria [21,26,30,37]. In the remaining twelve SRs, 50% to 74% of included studies fulfilled all the inclusion criteria [19,20,22–25,28,31,32,34–36].

All SRs primarily examined studies on school-based, rather than school-linked interventions. The 20 SRs covered eight health areas: nine on mental health [19,23,24,28,29,32–34,36], four on oral health [18,25,27,30], two on asthma [31,37], and one SR each on sleep [20]; obesity [26]; vision [21]; menstrual management [22]; and sexual and reproductive health (SRH) [35]. Eleven SRs included only cluster- and individually-randomized controlled trials [19–21,24,27,28,30,32,34,37], seven SRs included other types of controlled and uncontrolled experimental studies in addition to RCTs [22,25,26,31,33,35,36], and two SRs included only QEs [29] or controlled clinical trials [23]. Table 1 summarizes the characteristics of the 20 included SRs.

**Table 1.**
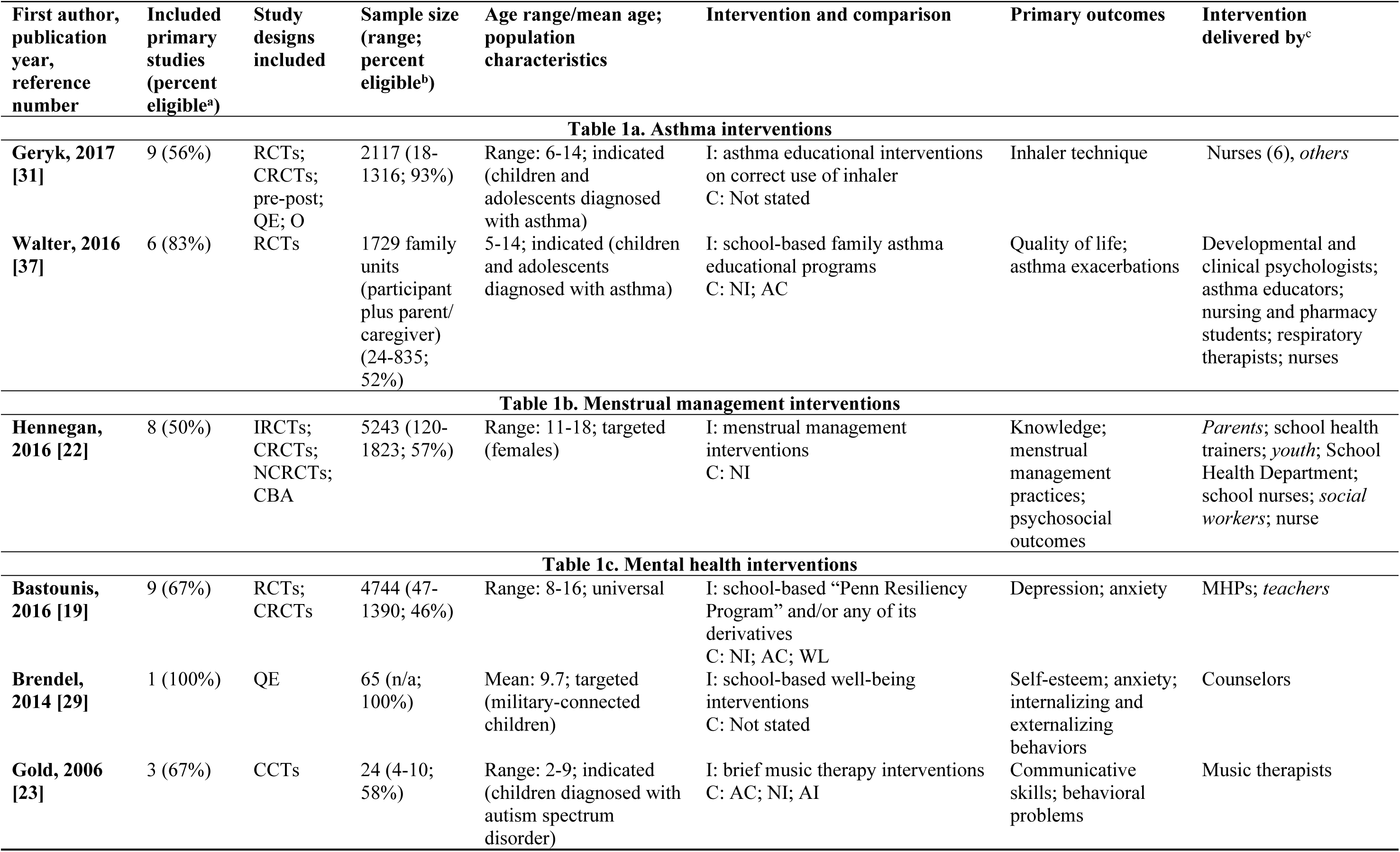
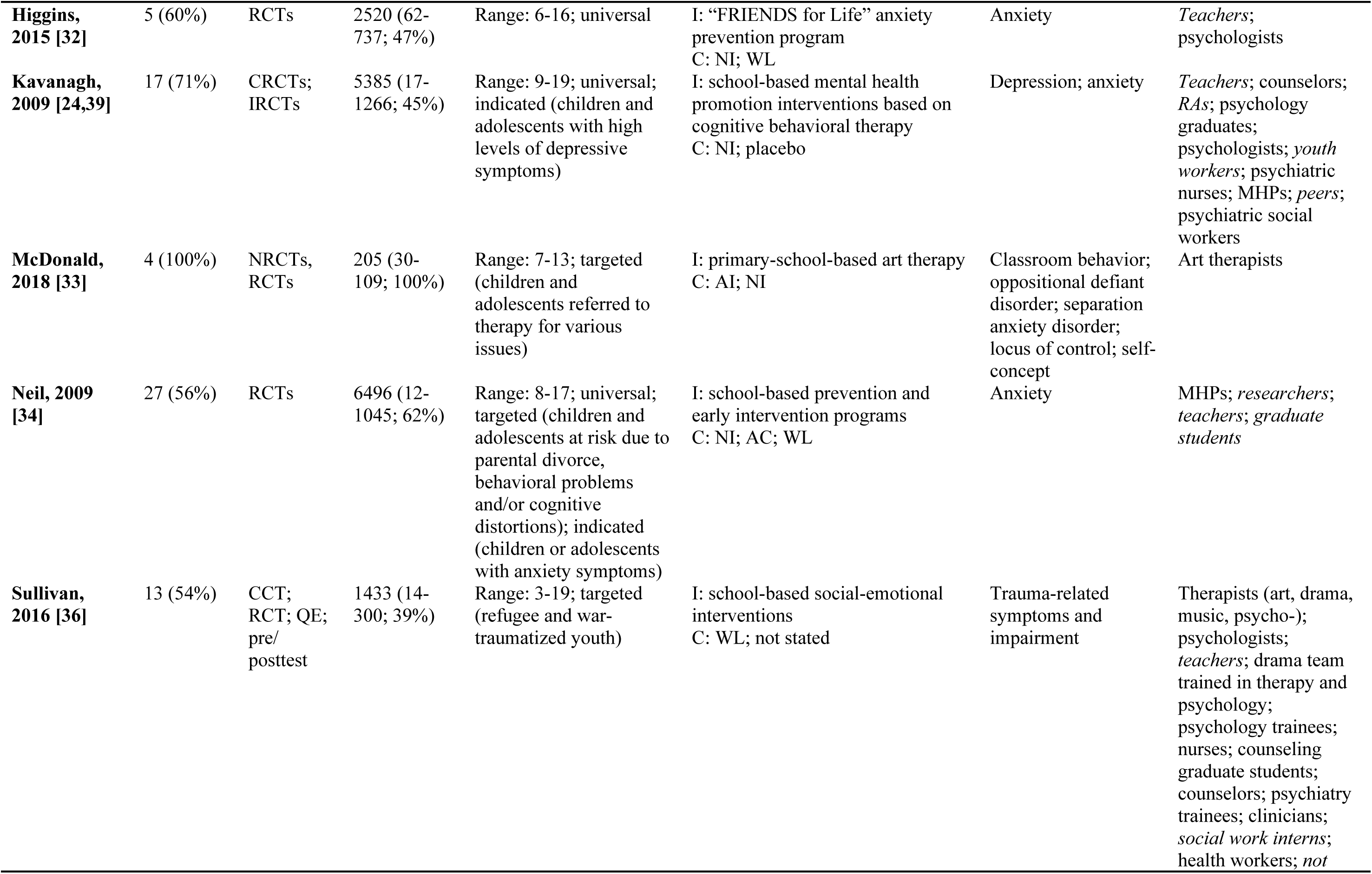
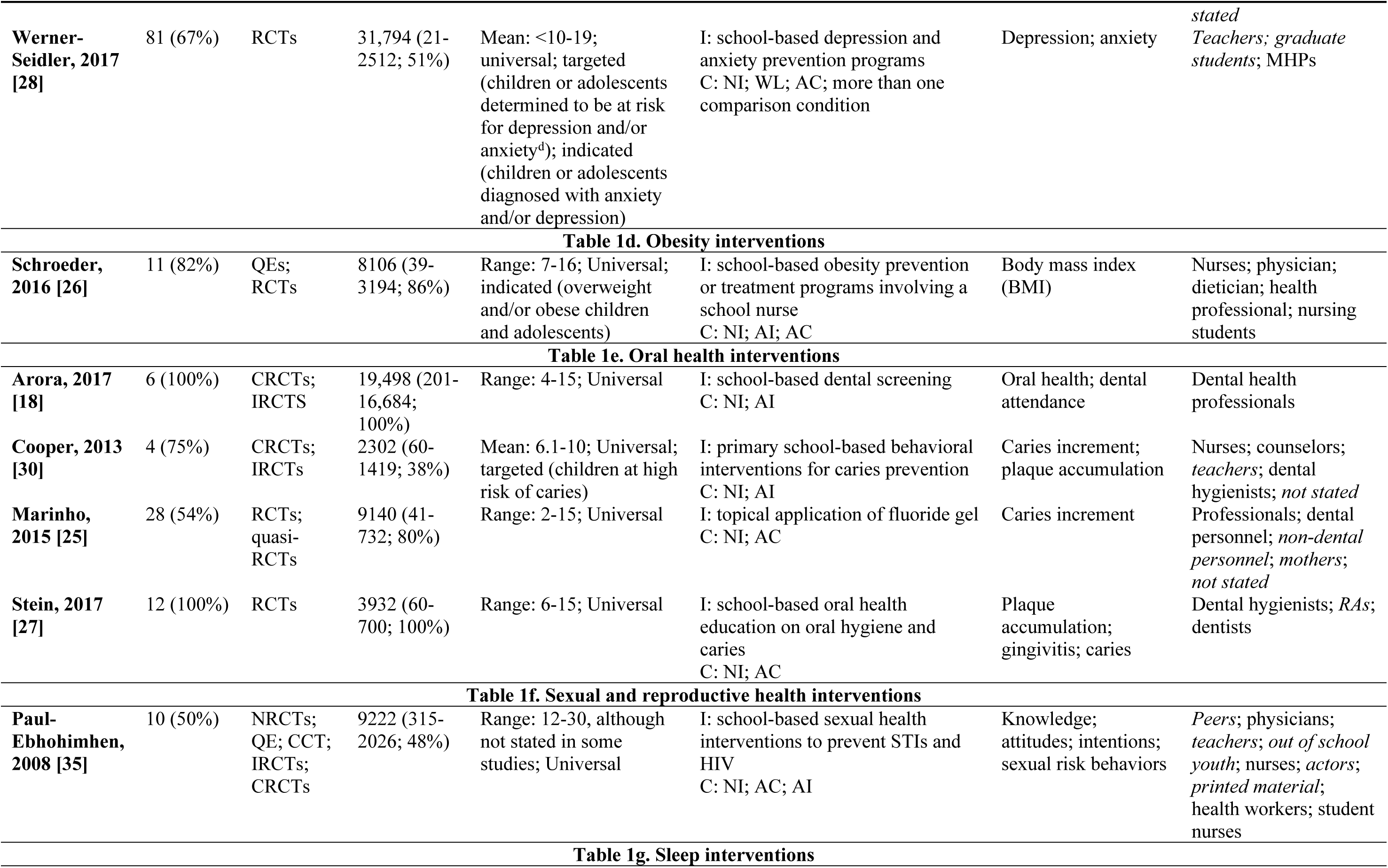
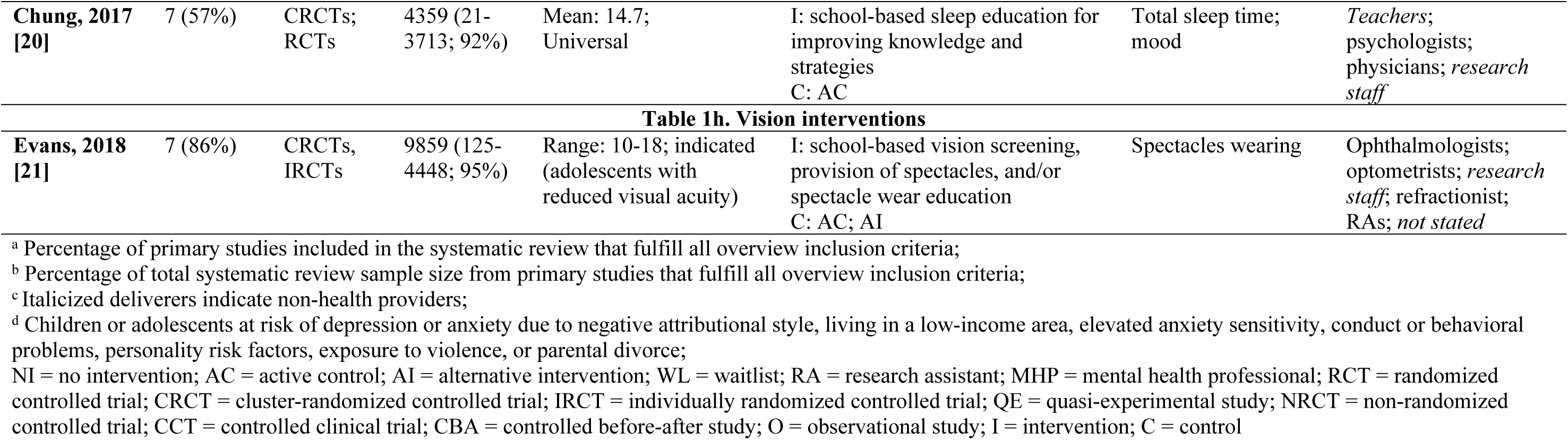
Characteristics of included systematic reviews

### Quality appraisal of systematic reviews within this overview

The corrected covered area (CCA) was found to be 1, indicating only slight overlap between the 20 SRs. Calculations for the CCA can be found in S5 Appendix. Table 2 presents the remaining three of the four items of Ballard and Montgomery’s checklist for overviews of reviews: (2) levels of confidence in results for each included SR, (3) publication year, and (4) match in scope to the overview. A majority of the studies (80%) were given low or critically low levels of confidence. Only three SRs [18,21,25] were scored as having moderate levels of confidence and just one [30] was given a high level of confidence. The details of the quality appraisal of primary studies included in the SRs are given in S4 Appendix.

**Table 2.** Items 2, 3 and 4 of Ballard and Montgomery’s four-item checklist for risk of bias in overviews of reviews^a^ [13]

| First author | Health area | AMSTAR 2 rating of confidence<sup>b</sup> (Item 2) | Published in 2016, 2017, or 2018? (Item 3) | At least 75% of included studies relevant<sup>c</sup>? (Item 4) |
| --- | --- | --- | --- | --- |
| Arora [18] | Oral health | Moderate | 2017 | 100% |
| Bastounis [19] | Mental health | Low | 2016 | 67% |
| Brendel [29] | Mental health | Critically low | 2014 | 100% |
| Chung [20] | Sleep | Low | 2017 | 57% |
| Cooper [30] | Oral health | High | 2013 | 75% |
| Evans [21] | Vision | Moderate | 2018 | 86% |
| Geryk [31] | Asthma | Critically low | 2017 | 56% |
| Gold [23] | Mental health | Low | 2006 | 67% |
| Hennegan [22] | Menstrual management | Low | 2016 | 50% |
| Higgins [32] | Mental health | Critically low | 2015 | 60% |
| Kavanagh [24,39] | Mental health | Critically low | 2009 | 71% |
| Marinho [25] | Oral health | Moderate | 2015 | 54% |
| McDonald [33] | Mental health | Low | 2018 | 100% |
| Neil [34] | Mental health | Critically low | 2009 | 56% |
| Paul-Ebhohimhen [35] | SRH | Critically low | 2008 | 50% |
| Schroeder [26] | Obesity | Critically low | 2016 | 82% |
| Stein [27] | Oral health | Low | 2017 | 100% |
| Sullivan [36] | Mental health | Critically low | 2016 | 54% |
| Walter [37] | Asthma | Low | 2016 | 83% |
| Werner-Seidler [28] | Mental health | Low | 2017 | 67% |

<sup>a</sup> The four items of the checklist include: overlap, AMSTAR 2 rating of confidence, up-to-date, and relevance of included studies
<sup>b</sup> See S6 Appendix for AMSTAR 2 rating information
<sup>c</sup> i.e., percentage of studies within the systematic review that clearly fulfill all inclusion criteria
SRH = sexual and reproductive health

### Findings: comprehensive, multi-component, or multi-health area services

None of the SRs evaluated comprehensive, multi-component, or multi-health area school health services.

### Findings: asthma interventions

Two SRs found strong evidence for the potential effectiveness of educational interventions for children and adolescents with asthma diagnoses (Table 3a) [31,37]. Geryk and colleagues found that education on correct use of an inhaler improved inhaler technique, regardless of deliverer, method, or duration of the intervention [31]. However, they did not assess risk of bias or appraise the quality of included studies [31]. Walter and colleagues found that family asthma educational programs for children and their parents or caregivers improved quality of life for both caregivers and children, and decreased asthma exacerbations for children [37]. While results from primary studies were statistically significant in both SRs, heterogeneity of interventions precluded meta-analysis by Walter and colleagues [37] and no reason was given for why meta-analysis was not performed by Geryk and colleagues [31].

**Table 3.**
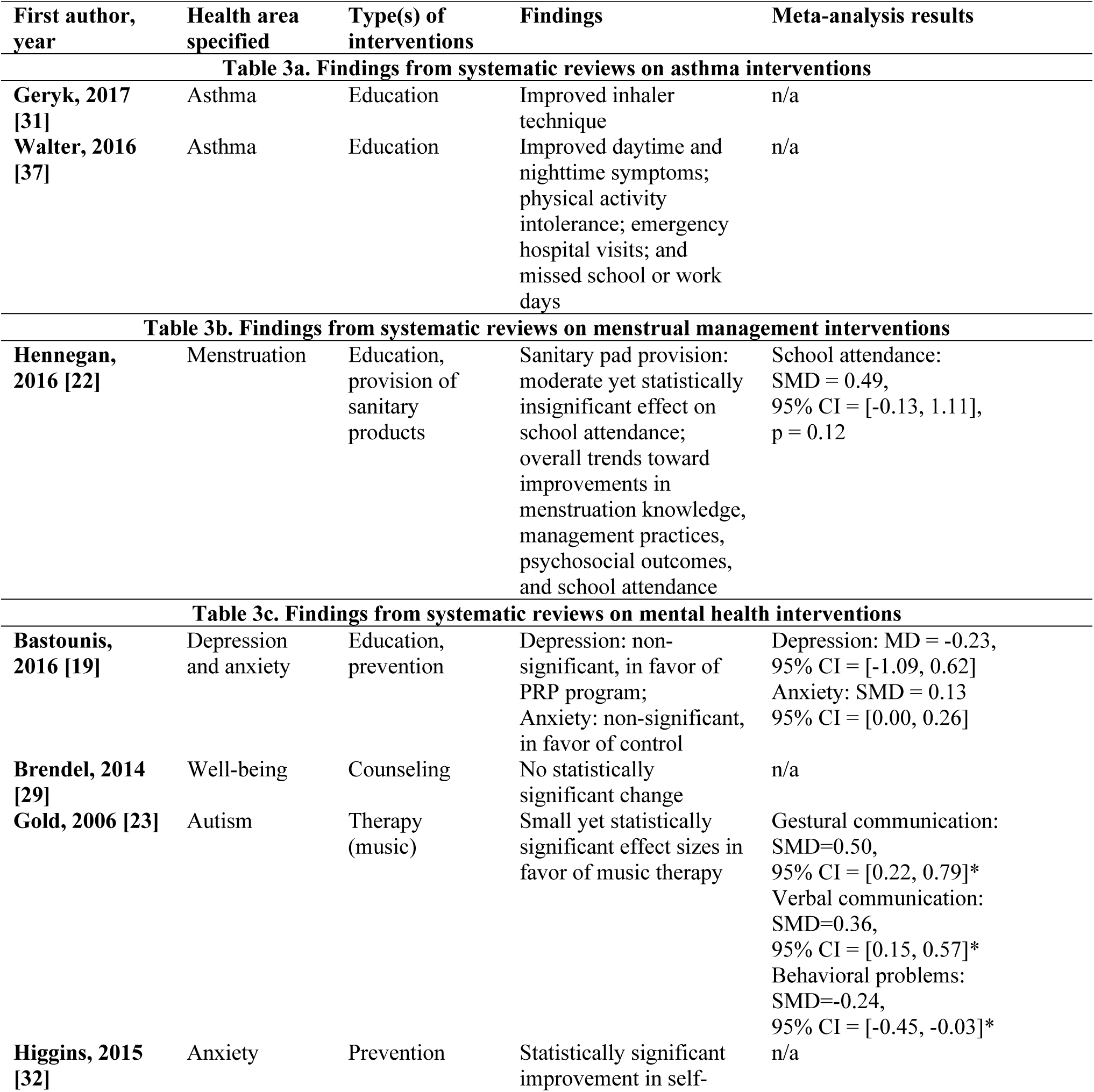
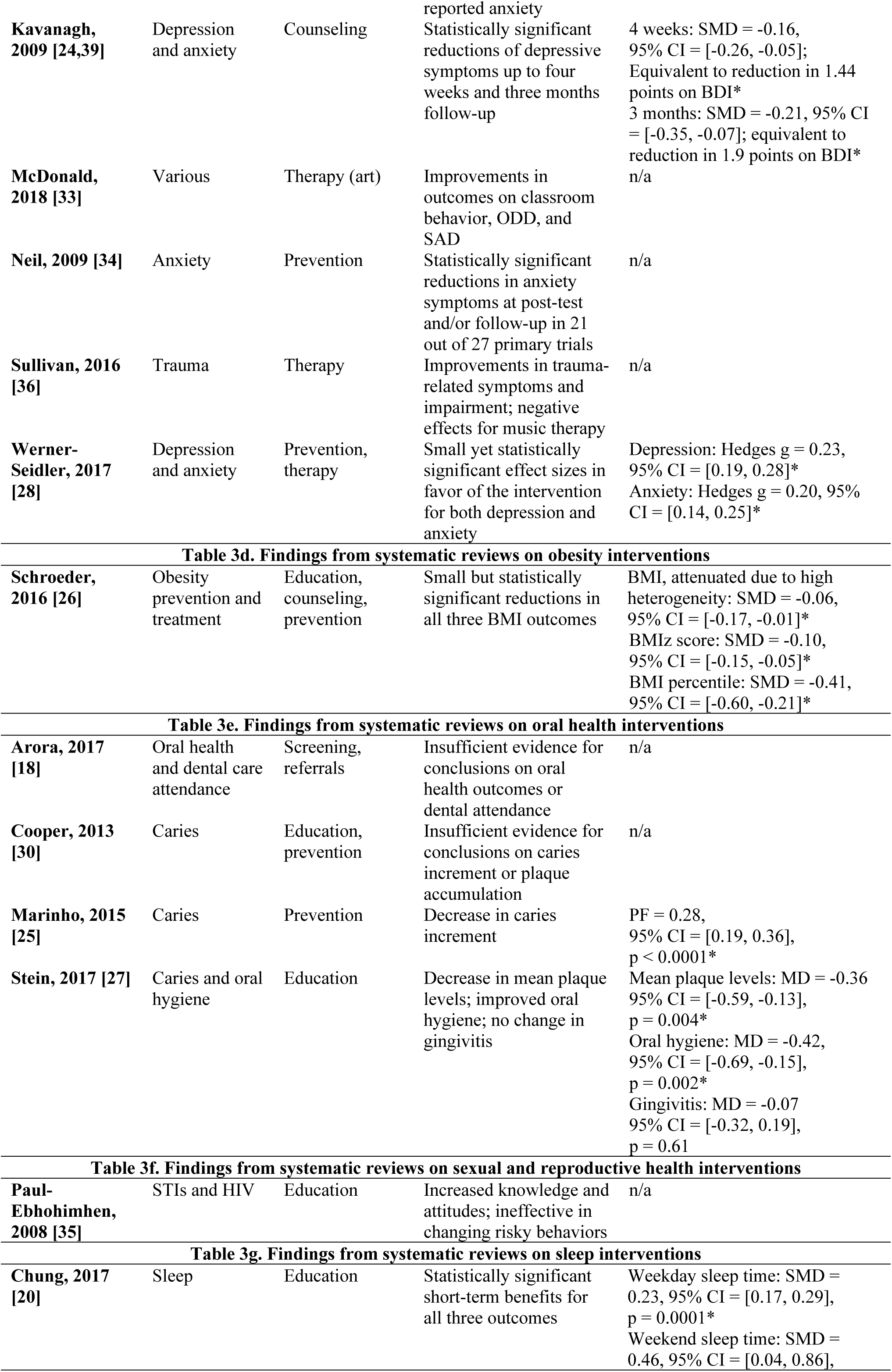
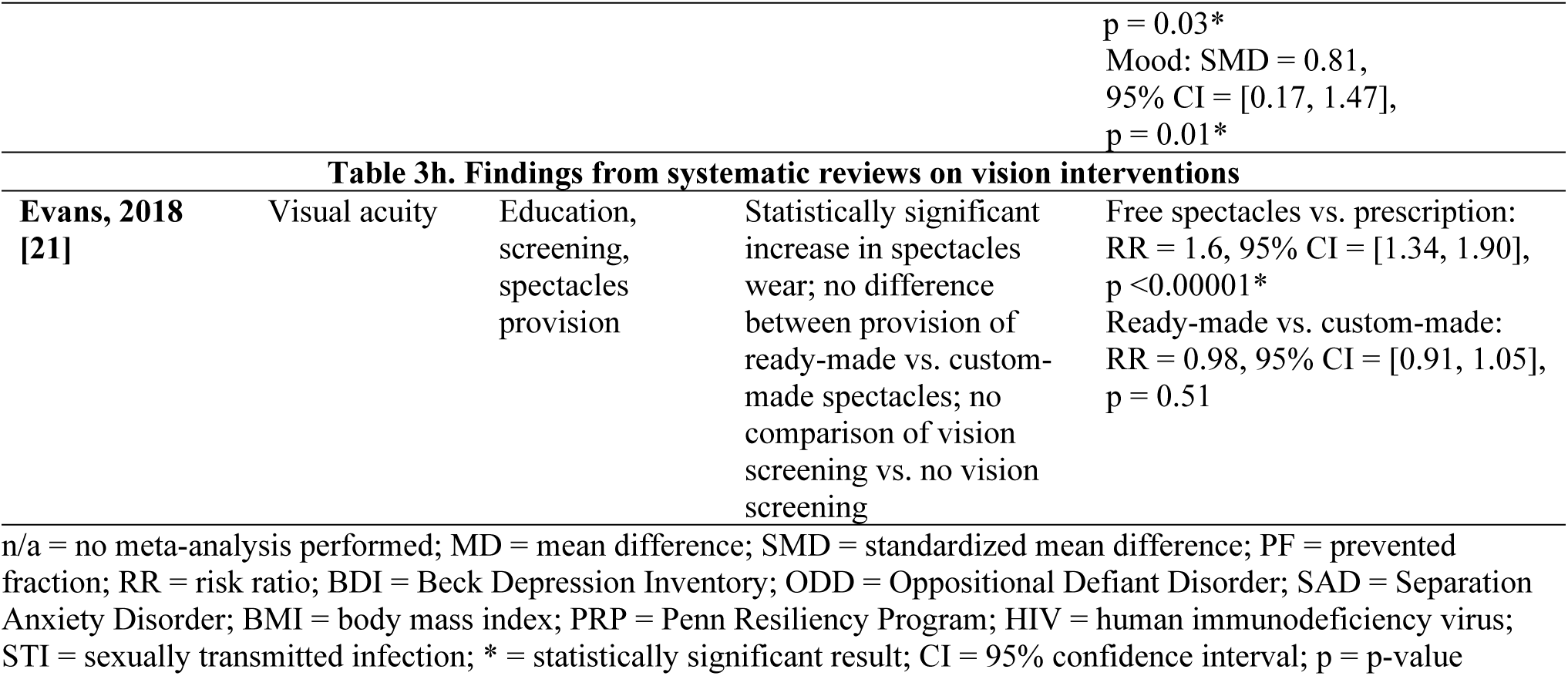
Findings from systematic reviews

### Findings: menstrual management interventions

Hennegan and Montgomery assessed the effectiveness of “hardware” and “software” menstrual management interventions (Table 3b) [22]. Hardware interventions included provision of sanitary products and software interventions focused on menstrual management education. A meta-analysis of two studies on sanitary pad provision found a moderate but statistically non-significant effect on school attendance. However, it was unclear whether these studies involved a health provider [22]. Outcomes across studies differed, but the authors noted trends toward improvement in menstruation knowledge, management practices, psychosocial outcomes, and school attendance. Hennegan and Montgomery found a high level of heterogeneity and substantial risk of bias in the included studies overall, thus they were unable to make conclusions about the effectiveness of menstrual management interventions [22].

### Findings: mental health interventions

The effectiveness of school-based mental health services was assessed in nine SRs (Table 3c) [19,23,24,28,29,32–34,36,39]. SRs addressed various intervention types: universal interventions [19,24,28,32,34,39]; targeted interventions for military-connected children [29], children and adolescents at risk for depression and/or anxiety [28,34], refugee and war-traumatized youth [36], and children referred to therapy [33]; and indicated interventions for children and/or adolescents diagnosed with autism spectrum disorder [23], depression [24,28,39], or anxiety [24,34,39].

Prevention and treatment of mood disorders was assessed in five SRs, all of which targeted children and adolescents using RCTs of established programs. Higgins and O’Sullivan assessed the FRIENDS for Life program, a manual-based cognitive behavioral anxiety prevention program comprised of ten sessions with developmentally-tailored programs for different age groups [32]. They found statistically significant improvements in self-reported measures of anxiety for participants who completed the program as compared to those in the control group [32]. A SR and meta-analysis by Bastounis and colleagues on the educational and preventative Penn Resiliency Program (PRP) and its derivatives found small, non-significant effect sizes for the prevalence of both depression and anxiety, favoring the intervention in the former and the control in the latter [19]. The remaining three SRs also assessed FRIENDS and PRP along with additional often-overlapping programs. Neil and Christensen analyzed 21 unique anxiety prevention and early intervention programs and found reductions in anxiety symptoms in 78% of the included studies [34]. Kavanagh and colleagues examined depression and anxiety group counseling programs based on cognitive behavioral therapy (CBT) and found statistically significant reductions of depressive symptoms at both four weeks and three months follow-up [24,39]. Finally, meta-analysis by Werner-Seidler and colleagues of 81 RCTs on the effectiveness of depression and anxiety prevention and group therapy programs found small yet statistically significant effect sizes in favor of the intervention groups for both depression and anxiety as compared to control groups [28]. Although the overall degree of overlap between all SRs within this overview was slight, the overlap between just these five SRs targeting mood disorders was high (CCA = 11).

Assessments of music [23] and art therapy [33] in two SRs reported weak evidence of effectiveness. Gold and colleagues assessed daily music therapy as an intervention to improve verbal and gestural communicative skills and reduce behavioral problems in children diagnosed with autism spectrum disorder [23]. Meta-analysis found small but statistically significant effect sizes in favor of music therapy for gestural communication, verbal communication, and behavioral problems [23]. McDonald and Drey narratively assessed group-based art therapy as an intervention for children with Oppositional Defiant Disorder (ODD), Separation Anxiety Disorder (SAD), moderate to severe behavior problems, or learning disorders [33]. The authors found improvements in classroom behavior, symptoms of ODD, and symptoms of SAD [33]. However, in the studies included in both Gold and colleagues’ and McDonald and Drey’s SRs, the numbers of participants per intervention group were very small: 4-10 and 12-25 per study, respectively, introducing possibility of bias [23,33].

Mostly favorable evidence of effectiveness was found in a SR of social-emotional interventions for refugee and war-traumatized youth from 26 countries [36]. Improvements in trauma-related symptoms and impairment were found through narrative assessment of creative expression interventions, cognitive behavioral interventions, and multifaceted interventions [36]. In contrast with Gold and colleagues [23], this SR by Sullivan and Simonson found negative effects from music therapy interventions [36]. However, there was no risk of bias assessment in this SR and therefore the results must be interpreted cautiously.

The final SR on mental health services examined well-being interventions for children with a parent in the military [29]. Only one quasi-experimental study from the United States in 1999 was included in the SR. The study assessed a group counseling intervention and found no statistically significant effects on the prevalence of anxiety, self-esteem, internalizing behavior or externalizing behavior [29].

### Findings: obesity interventions

Schroeder and colleagues reviewed the effectiveness of obesity treatment and prevention interventions that specifically involved a school nurse (Table 3d) [26]. Most interventions involved school-nurse-delivered nutrition counseling, nutrition and health education, and some parent involvement or physical activity. Meta-analysis indicated small, yet statistically significant, reductions in body mass index (BMI), BMI z-score, and BMI percentile for both obesity treatment and prevention [26].

### Findings: oral health interventions

SRs on oral health interventions focused on prevention [25,30], screening [18], and education [27,30] (Table 3e). Strongest evidence in favor of oral health interventions emerged from a SR on universal topical application of fluoride gel for the prevention of dental caries [25]. Meta-analysis results indicated a statistically significant effect on the before-after change in caries prevalence [25]. A universal educational intervention on oral hygiene and caries produced weaker evidence of effectiveness [27]. Small but statistically significant effect sizes were found in favor of the intervention for mean plaque levels and oral hygiene, but no statistical significance was found for change in gingivitis indices [27].

Two SRs on dental health screening [18] and behavioral interventions for caries prevention [30] found limited evidence of effectiveness. Arora and colleagues did not find any RCTs that looked at the effectiveness of dental health screening versus no screening on improving oral health outcomes, but their search did locate six RCTs from the United Kingdom and India with dental care attendance as the outcome [18]. The data was too heterogeneous to meta-analyze, and the authors determined that the certainty of the evidence of the benefit of dental screening in increasing dental attendance was very low [18]. The other SR examined behavioral interventions in the form of education on tooth brushing and the use of fluoride toothpaste in Brazil, Italy, United Kingdom, and Iran [30]. Due to the diversity in outcome measures and intervention intensities, the authors felt unable to make any evidence-based recommendations [30].

### Findings: sexual and reproductive health interventions

A SR on sexual health interventions for prevention of sexually transmitted infections (STIs) and human immunodeficiency virus (HIV) in sub-Saharan African countries found that educational interventions were successful in increasing knowledge and attitudes for participants (Table 3f) [35]. However, the SR suggested that the studies were ineffective in changing self-reported risky behaviors, although follow-up was either immediate or short-term (less than 6 months) [35]. This SR did not discuss the quality or risk of bias of included studies [35].

### Findings: sleep interventions

Chung and colleagues systematically reviewed and meta-analyzed universal sleep education programs as compared to no additional sleep intervention from Australia, New Zealand, Brazil, and Hong Kong (Table 3g) [20]. Five of the included studies examined the same weekly sleep education program from the Australian Centre for Education in Sleep. The sixth study assessed a 4-day program in Brazil. Meta-analysis of the six studies showed statistically significant short-term benefits for weekday sleep time, weekend sleep time, and mood [20]. However, these results did not persist at follow-up [20].

### Findings: vision interventions

Evans and colleagues reviewed seven RCTs from China, India, and Tanzania on vision screening for correctable visual acuity deficits at or before school entry (Table 3h) [21]. Through meta-analysis of two RCTs, the authors found that school vision screening combined with provision of free spectacles resulted in a statistically significant 60% increase in the wearing of spectacles at 3-8 months follow-up as compared to vision screening combined with prescription for spectacles only [21]. Evans and colleagues found no statistically significant difference in the proportion of students wearing spectacles at 1-4 months follow-up between vision screening with provision of ready-made spectacles and vision screening with provision of custom-made spectacles in a meta-analysis of three RCTs [21]. Education on the wearing of spectacles in addition to vision screening as compared to vision screening alone did not have a significant effect [21]. No SRs found eligible studies comparing vision screening with no vision screening.

## DISCUSSION

### Summary of key findings

This overview found 20 SRs covering 270 primary studies. The majority of SRs assessed educational, counseling, or preventive interventions, most of which were special research interventions rather than routinely-delivered school health services. No SR examined comprehensive or multi-component school health services, despite the fact that comprehensive services may be more efficient, easier to implement, and more sustainable than single interventions [40]. Results from this overview suggest that certain interventions can be effective in improving child and adolescent health outcomes, and thus may be worthwhile for integration into school health programs.

### Effectiveness of specific interventions

Vision screening is one of the most common forms of school health services [9], although the majority of programs are concentrated in high-income countries (HIC) [21]. Although prevalence of visual impairment varies widely by ethnic group and age [41], WHO estimates that at least 19 million children below age 15 are visually impaired [42]. Evans and colleagues found strong evidence from China, Tanzania, and India that school vision screening for correctable visual acuity deficits increased wearing of spectacles when spectacles were provided at no cost [21]. A recent guideline from the International Agency for Prevention of Blindness (IAPB) reiterates the importance of free spectacles and goes further to suggest that low- and middle-income countries (LMIC) adopt comprehensive school eye health programs [43]. Vision screening linked with free provision of spectacles, as a component of a comprehensive school eye health program, is an example of a cost-effective form of school health services that may be implemented.

Five SRs covered depression and/or anxiety prevention and early intervention programs, with the FRIENDS for Life program (FRIENDS) and the Penn Resiliency Program (PRP) most common [19,24,28,32,34]. Given that FRIENDS has been endorsed by WHO [44] and was found to be effective in decreasing anxiety symptoms in all four SRs where it was mentioned in this overview [28,32,34,39], policy makers and school health officials may consider incorporating this or similar programs into existing school health services. The four SRs that included PRP found mixed evidence [34] or no evidence of effectiveness [19,24,28,39], bringing the popularity of this intervention into question. Finally, creative therapy interventions seem to be effective for indicated populations of school-age children, such as children with autism spectrum disorder [23,33]. However, this conclusion should be interpreted cautiously due to small effect sizes, small sample sizes, and conflicting evidence on the effectiveness of music therapy between Sullivan and Simonson [36] and Gold and colleagues [23].

Comprehensive school programs that promote healthy school environments, health and nutrition literacy, and physical activity are one of the six key areas for ending childhood obesity recommended by WHO [45]. This overview found only one SR that assessed obesity treatment and prevention delivered by a health professional in schools, despite the fact that over 340 million school-age children and adolescents were overweight or obese in 2016 [46]. Schroeder and colleagues found that school nurses are well positioned to deliver nutritional counseling, design and coordinate physical activity interventions, and educate parents, students, and staff on health, nutrition, and fitness [26]. However, all included primary studies were delivered in HIC [26].

Schools are considered to be an ideal platform for oral health promotion through education, services, and the school environment [6]. The most promising evidence from a SR was on topical application of fluoride gel for the prevention of dental caries [25]. Educational interventions had mixed effects. A SR that focused on behavioral education, such as demonstrating how to correctly brush teeth, found no evidence for reduction in caries [30], whereas a SR on oral hygiene and caries education found evidence for decreased plaque and improved hygiene [27]. More research should be done to identify the content and methods of deliver that make some oral health education interventions more effective than others.

### Overall effectiveness of school health services

It is difficult to determine overall effectiveness of school health services from this overview because the included SRs do not sufficiently cover the health areas most relevant for children and adolescents. In 2015, the top five leading causes of death for 5-9 year olds were lower respiratory infections, diarrheal diseases, meningitis, drowning, and road injury [47]. Among 10-14 year olds, the leading causes of death were lower respiratory infections, drowning, road injury, diarrheal diseases, and meningitis [47]. Finally, for 15-19 year olds, the leading causes of death were road injury, self-harm, interpersonal violence, diarrheal diseases, and lower respiratory infections [47]. Leading causes of disability-adjusted life years (DALYs) for 10-19 year olds were iron deficiency anemia, road injury, depressive disorders, lower respiratory infections, and diarrheal diseases [47]. This overview shows that the current SR literature does address mental health, specifically mood disorders. However, the causes of death and disability beyond self-harm and depressive disorders are currently not addressed. Although mortality and morbidity statistics vary by region and country, it is clear that the health areas included in this overview reflect a small subset of the global burden of disease for children and adolescents.

Furthermore, this overview exposes a mismatch between the SR literature on effectiveness of school health services and the actual school health services that are most commonly delivered. Vaccinations have been identified as the most common type of intervention in schools in at least 35 countries or territories [9], and there is evidence of effectiveness from primary studies regarding feasibility of school-based vaccination programs [48,49]. Yet no SRs on vaccinations fulfilled the inclusion criteria for this overview, suggesting the need for these SRs to be conducted. Additionally, at least 94 countries or territories include some form of school health services that are routinely delivered, as opposed to special research interventions [9]. This overview primarily found evidence for special research interventions, suggesting a need for assessment of routinely-delivered school health services. One of the central questions of this overview was whether the school health services that are regularly delivered across the globe are evidence-based. The mismatch in the SR literature identified by this overview demonstrates that more research must be done before an answer to this question can be determined.

Another important gap that this overview reveals is a lack of research on interventions carried out in LMIC and low-income countries (LIC). Only one of the 20 SRs included in this overview examined studies from a majority of LMIC and LIC [22,38]. This is problematic given that health disparities for children and adolescents are greater in LMIC and LIC than in higher income countries [10]. Additionally, resources differ by income level and therefore effective interventions in HIC may need to be tailored or changed entirely in order to be feasible in LMIC and LIC. WHO reports densities of less than one physician per 1000 population in 76 countries and less than three nurses or midwives per 1000 population in 87 countries [50]. Thus, responsibility for interventions may need to be redistributed in countries with limited access to health professionals. This may be solved by coordinating school visits from hospital-based health professionals or by linking schools with local health centers for children and adolescents to access as needed. Research on the effectiveness of school health services in LMIC and LIC must be prioritized in order to fulfill Sustainable Development Goal 3.8 and reach universal health coverage by 2030 [51].

Although three SRs mentioned the cost of specialized professionals delivering interventions versus teachers or a school nurse [26,32,36], cost, let alone cost-effectiveness, was not closely analyzed in any of the included SRs. For useful recommendations to be made regarding school health services, cost-effectiveness must be more closely examined by primary studies and SRs.

### Methodological limitations of overviews of systematic reviews

Although overviews offer a comprehensive method for synthesizing evidence, they also come with important methodological limitations. First, an overview is unlikely to include the latest evidence if recent primary studies have not yet been included in SRs. This lag may preclude the ability for an overview to truly reflect current knowledge [13]. While this overview found significant gaps in the evidence for certain health areas, this does not necessarily mean that relevant high quality primary trials have not been conducted.

Second, the ability of overviews to make valid and accurate conclusions is dependent upon the accuracy, rigor, and inclusiveness of the SRs themselves. 80% (n=16) of SRs included in this overview were given ratings of low or critically low confidence using the AMSTAR 2 checklist, although this is not unusual given the stringency of the checklist. Nonetheless, it is interesting to note that all four of the SRs given moderate or high levels of confidence were Cochrane reviews. The remaining Cochrane review was given a critically low level of confidence, though this may be because it was published in 2006 and standards for both the methods and reporting of Cochrane reviews have improved in recent years.

Third, the scopes of individual SRs often differ from the scope of the overview, a problem that Ballard and Montgomery call a “scope mismatch” [13]. In this overview, SRs with at least 50% of included studies fulfilling all criteria were included after extensive discussion between the authors and experts in the field. This implies that a narrower range of SRs would have been eligible if a stricter cut-off had been selected, and vice versa. It is important to take this into account when interpreting results.

Finally, overlap of primary trials between SRs can bias results of an overview [15]. There is no definitive guidance on how to correct for overlap, as both including or excluding overlapping SRs presents potentially biased results [13,15]. This overview measured overlap using the corrected covered area (CCA) [16] and did not exclude overlapping studies. However, the degree of overlap across all 20 SRs was graded as being small (CCA = 1), while there was high overlap in SRs on mood disorders (CCA = 11). CCA values for all health areas and calculations are available in S5 Appendix.

### Additional strengths and limitations of this overview

A key limitation of this overview is that only publications self-titled “systematic reviews” were included. This decision was made because of the vast numbers of reviews available and the increased rigor associated with the term “systematic”. A sensitivity analysis comparing “systematic*” with “systematic review” found that the number of search results increased almost three-fold, but did not reveal any new articles that would eventually have met the subsequent eligibility criteria.

Another limitation is that this overview only included randomized and non-randomized controlled trials, quasi-experimental studies and other controlled study designs where health professionals delivered the intervention. While this strengthens the rigor of included studies and improves decision-making ability, it also excludes potentially relevant literature.

A strength of this overview is that it attempted to answer a question that has not yet been answered regarding the effectiveness of both comprehensive and specific school health services delivered by a health provider. While other pillars of Health Promoting Schools have relevant guidance documents, guidance on school health services is limited and not explicitly evidence-based. Given the wide reach of schools and the fact that school health services already exist in most countries, international guidelines are needed to clarify whether school health services can be effective, and if so, which interventions should and should not be included. This overview makes an important first step toward that guideline.

## CONCLUSIONS

This overview presents multiple effective interventions that may be offered as a part of school health services delivered by a health provider. However, it is difficult to formulate an overarching answer about the effectiveness of school health services for improving the health of school-age children and adolescents due to the heterogeneity of SRs found and the evident gaps in the SR literature. More than half of included SRs analyzed mental health and oral health interventions, and no SRs were found that assessed other relevant health areas, such as vaccinations, communicable diseases, injuries, etc. Further, no SRs evaluated comprehensive or multi-component school health services. If school health services are to truly improve the health of children and adolescents, they must comprehensively address the most pressing problems of this population. In order for policy makers and leaders in school health to make evidence-based recommendations on which services should be available in schools, who should deliver them, and how should they be delivered, more SRs must be done. These SRs must assess routine, multi-component school health services and the characteristics that make them effective, with special attention to content, quality, intensity, method of delivery, and cost. The gaps in the SR literature identified by this overview will inform the commissioning of new SRs by WHO to feed into evidence-based global recommendations.

## ACKNOWLEDGEMENTS

We would like to thank Tomas Allen for his guidance and support in designing the search strategy. We are also grateful for advice from the WHO school health services guideline Steering Group and Guideline Development Group members.

## SUPPORTING INFORMATION CAPTIONS

**S1 Appendix.** Protocol

**S2 Appendix.** Search strategy

**S3 Appendix.** Excluded full text articles

**S1 Table**. Excluded full text articles and reasons for exclusion

**S4 Appendix**. Quality appraisal of primary studies

**S2 Table**. Quality appraisal of primary studies within included systematic reviews

**S5 Appendix**. Corrected covered area

**S3 Table**. Calculation of corrected covered areas (CCAs)

**S4 Table**. Classification of primary studies within included systematic reviews for use in corrected covered area calculation

**S6 Appendix**. AMSTAR 2 classifications and results

**S5 Table**. AMSTAR 2 checklist with designations of critical and non-critical

**S6 Table**. Answers to AMSTAR 2 checklist questions 1-16

**S7 Appendix**. PRISMA checklist

**S8 Appendix references**

